# Mouse Models Uniformly Featuring Human-like Lesions Harboring Drug-tolerant *Mycobacterium tuberculosis*

**DOI:** 10.1101/2025.10.17.683099

**Authors:** Gaëlle Guiewi Makafe, Tim Low-Beer, Kelsey Travis, Laura Cole, Derek Bernacki, Debra Duso, Matthew Cole, Larry Kummer, Alan Roberts, Tricia Hart, Kathryn Roemer, William Reiley, Mike Tighe, Brian Weinrick

**Affiliations:** Trudeau Institute, Saranac Lake, New York, USA

## Abstract

Mouse models have been key to studies of tuberculosis pathogenesis and drug efficacy, but many, such as those employing BALB/c mice, fail to reproduce the full range of heterogenous microniches observed in well-structured human lesions, which feature hypoxic caseous cores of necrotic debris surrounded by infected foamy macrophages. The granuloma presents a variety of environments differing in levels of oxygen, ions, nutrients, and intra versus extracellular residence, which determine the physiological state of the infecting bacillus and its susceptibility to immune or drug control. Recently, alternative mouse strains such as C3HeB/FeJ have allowed the study of infection and treatment in the context of these varied environments but exhibit substantial inconsistency in development of human-like lesions, both within and between individual mice. Building on the observation that inducible nitric oxide synthase (*Nos2*)-deficient mice consistently develop hypoxic necrotic lesions, we have established two simplified models with infection by the aerosol route. The first uses the slightly attenuated *M. tuberculosis* R1Rv strain, which produces a progressive infection that is contained at a high stable burden by an adaptive immune response. In the second model, vaccination with the attenuated Δ*RD1*, pantothenate auxotroph mc^2^ 6230 protects from an otherwise lethal infection with virulent *M. tuberculosis* Erdman. This model reflects most contemporary tuberculosis infections, which take place in the context of a pre-existing immune response from vaccination. Both variations uniformly develop well-structured hypoxic necrotic lesions harboring drug tolerant bacteria. These refined models will be useful in studies of *M. tuberculosis* infection and treatment.

## Introduction

Evaluation of the efficacy of experimental tuberculosis therapeutics is commonly performed using mouse models, due to the relatively small size and low cost of mice. The models used most frequently involve infection of C57BL/6 or BALB/c mice with approximately 100 colony forming units (CFU) of *Mycobacterium tuberculosis* (*Mtb or M. tuberculosis*) by aerosol inhalation [1]. In many cases, these traditional models effectively mimic the responses of human tuberculosis patients to chemotherapy and allow an accurate prediction of the outcomes of clinical trials.

### Limitations of current tuberculosis drug efficacy mouse models

Occasionally, however, tuberculosis mouse models have come up short in predicting drug efficacy in humans. Although studies using BALB/c mice suggested that substitution of a fluoroquinolone into standard combination therapy for ethambutol or isoniazid could allow the shortening of treatment from six months to four months [2], the REMoxTB clinical trial demonstrated that these shorter regimens were inferior to the six-month standard of care (SOC) regimen in achieving relapse-free cure [3].

A central hypothesis concerning the failure of certain mouse models to accurately predict human outcomes suggests that it is the inability of these models to reproduce important aspects of tuberculosis pathology present in human infections that limits their predictive power [4]. Specifically, a key feature of human tuberculosis pathology is the formation of well-structured necrotic granulomas. These lesions consist of a hypoxic caseous necrotic core surrounded by a rim of infected macrophages, further enclosed within a layer of lymphocytes and fibroblasts [5]. This type of pathology is not present in most commonly used mouse models, which instead show a diffuse inflammatory cellular infiltration, in which the bacteria are almost exclusively within phagocytes [6]. The granuloma presents a variety of distinct microenvironments to the infecting bacteria, many of which are extracellular, and could allow more drug-tolerant differentiated states [7]. Additionally, the caseous core has been shown to accumulate reduced concentrations of some drugs such as fluoroquinolones, further limiting potential efficacy [8].

### Caseous hypoxic granulomas in mice

There are a few tuberculosis mouse models described that do generate structured granulomas, the most widely used employs the C3HeB/FeJ mouse [9]. The effectiveness of moxifloxacin-containing regimens in this model for treatment-shortening more closely reproduced clinical trial results, with a one-month reduction in time to relapse-free cure seen when moxifloxacin was substituted for isoniazid, versus the two-month reduction seen in a BALB/c model [10]. The C3HeB/FeJ model has several limitations, however. The mouse strain is very susceptible to tuberculosis; even when the infecting dose is kept within a tight window of 50-80 CFU it is common for 20% of C3HeB/FeJ mice to succumb to the disease before the development of granulomas. Additionally, a subset of mice does not generate granulomas, and within those that do there is often a mix of granulomas and the diffuse cellular inflammatory infiltrates [11]. This heterogeneity in pathology can lead to heterogeneity in drug responses, in some cases resulting in a curious bimodal distribution of drug efficacy [12, 13]. A model lacking this heterogeneity would be very valuable for the preclinical evaluation of novel tuberculosis chemotherapeutic regimens.

Fortunately, a novel mouse model that consistently develops necrotic, hypoxic, human-like granulomas employing iNOS-deficient mice was described by Reece and colleagues in the course of their investigation of immune mechanisms under hypoxia in *Mtb* infection [14]. Importantly, this model accurately predicts the differential activity of specific drugs in humans [15]. Unfortunately, the model is rather complex, and infection is not by the natural route. In brief, the model consists of intradermal infection with 10^4^ CFU in the ear, followed by TNF-α neutralization at two and three-weeks post-infection. By eight-weeks post-infection, the bacterial burden in the lungs reaches 10^6^ CFU and the characteristic human-like pathology begins to develop. At that point drug treatment responses observed reflect those seen in treatment of humans. The requirement for the intradermal infection followed by TNF-α neutralization to induce dissemination to the lungs is because mice lacking *Nos2* are unable to control an aerosol infection with *Mtb* type strain H37Rv and rapidly succumb. Interestingly, early studies of *Mtb* infection in these *Nos2^tm1Lau^* homozygous mice revealed that aerosol infection with a partially attenuated strain, R1Rv, results in progressive acute infection that is controlled but not cleared by an adaptive immune response, establishing a high-burden chronic infection [16].

*Mycobacterium tuberculosis* R1Rv, derived from the “R1” strain isolated in 1891 from a miliary tuberculosis patient, was first passaged in a rabbit and eventually demonstrated reduced virulence in guinea pigs [17, 18]. Whole genome sequencing (deposited as ATCC 35818 / TMC#205) revealed that R1Rv is closely related to H37Rv. Notable genetic variants that may contribute to its attenuated phenotype include frameshift mutations in phosphate-specific transport genes (*pstA1*, *pstS3*) and a gene required for sulfolipid biosynthesis (*pks2*).

In this study, we aimed to reproduce the key pathological features of the *Nos2-* deficient model while simplifying the experimental design by initiating infection through the natural aerosol route. We established and compared disease progression and granuloma pathology in two models: one using aerosol infection with the partially attenuated *Mtb* R1Rv strain, and the other involving prior vaccination with attenuated *Mtb* mc² 6230 followed by aerosol challenge with virulent *Mtb* Erdman. Additionally, we assessed the therapeutic responsiveness of the vaccinated model to monotherapy with multiple anti-tubercular drugs, providing insights into how lesion architecture influences treatment efficacy.

## Results

### The R1Rv Strain of *M. tuberculosis* Establishes Progressive, High-Burden Infection in *Nos2-*deficient Mice featuring Necrotic Granulomas

In this initial model, we sought to replicate findings from a study which demonstrated that *Nos2*-deficient mice could control an aerosol infection with the *M. tuberculosis* R1Rv strain over a 60-day period [19]. Notably, lung lesions in these mice resembled the well-structured granulomas typically observed in human tuberculosis [20].

Seeking to achieve a high burden chronic infection suitable for drug efficacy studies *Nos2*-deficient mice were aerosol-infected with approximately 10³ CFU of *Mtb* R1Rv. Lung bacterial burdens increased steadily over time, reaching a high and stable plateau of approximately 10⁶ CFU by week 6 in both female and male mice (Fig. 1A). This burden was maintained and slowly increased to around 10⁷ CFU by week 20 post-infection (data not shown).

**Figure 1.**
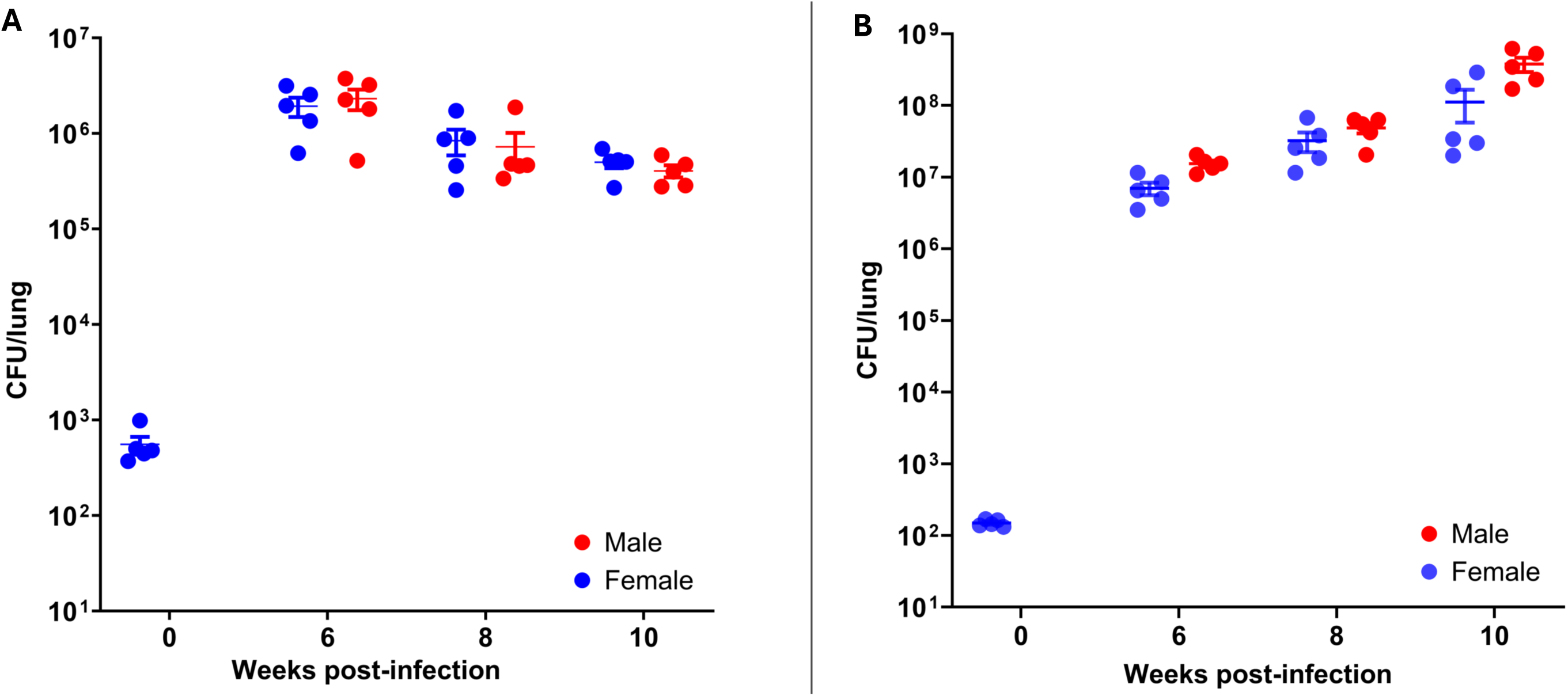
Bacterial burden kinetics in the lungs of *Nos2*-deficient mice following aerosol infection. Viable bacterial counts in the lungs of male (red circles) and female (blue circles) *Nos2*-deficient mice harvested at weeks 0, 6, 8, and 10 post-infection. **(A)** Naïve model: *Nos2*-deficient mice challenged with *Mtb* R1Rv. **(B)** Vaccinated model: *Nos2*-deficient mice previously vaccinated with mc^²^6230 and then challenged with *Mtb* Erdman-pYUB1785. Each data point represents an individual mouse (n = 5 per time point), with group means ± s.e.m. also shown.

This persistent infection was accompanied by progressive lung pathology, characterized by the development of well-structured granuloma lesions, which became increasingly pronounced between week 8 and week 20 post-infection. Thin-cut sections (5 μm) of fixed, paraffin-embedded left lung lobes were stained with hematoxylin and eosin (H&E) (Supplementary Fig. 1) or processed for immunohistochemistry to visualize hypoxic regions (Supplementary Fig. 2).

Importantly, the absence of significant mortality in this model supports its utility as a robust platform for studying chronic *Mtb* infection and for evaluating anti-tubercular candidates in the context of advanced, human-like pathology.

### Consistent Formation of Hypoxic Necrotic Granuloma and High Lung Burden in *Nos2*-deficient Mice Vaccinated with *mc² 6230* and Aerosol Challenged with *Mtb* Erdman

Whereas *Nos2* is not required for controlling infection with *Mtb* R1Rv, it is required to control virulent *Mtb* strain H37Rv, as *Nos2*-deficient mice succumbed within 40 days [19]. Similarly, MacMicking et al. (1997) reported rapid mortality (33–45 days post-infection) in *Nos2*-deficient mice intravenously infected with virulent *Mtb* Erdman [21]. In contrast, other investigators have observed control of virulent *Mtb* without *Nos2* [22]. This raises the possibility that immunization may allow *Nos2*-deficient mice to control infection with virulent *Mtb*. We hypothesized that vaccination-induced protective immunity might prolong survival and alter disease progression in *Nos2*-deficient mice.

In our initial protection assay, subcutaneous administration of ∼10⁷ CFU of BCG Pasteur, followed by aerosol challenge with ∼100 CFU *Mtb* H37Rv resulted in lung bacterial burdens and necrotic lesions comparable to those observed in the naïve R1Rv model (data not shown). Visible lesions were also noted at the injection site, indicating failure to limit the growth of the vaccine strain. As an alternative, we used *Mtb* mc²6230, in which RD1, the primary attenuating deletion in BCG, has been deleted, as well as *panCD*, which are required for pantothenate biosynthesis. This strain has been shown to stimulate robust immune responses but is unable to grow even in highly susceptible hosts such as SCID or CD4-deficient mice [23]. *Nos2*-deficient mice were vaccinated subcutaneously with ∼10⁷ CFU of mc² 6230, then aerosol-challenged 4 weeks later with ∼100 CFU of *Mtb* Erdman-pYUB1785, which bears a dual fluorescent reporter construct for the identification of persister cells [24]. The use of this reporter strain was to enable the spatial and temporal characterization of drug-tolerant *Mtb* cell populations during infection and treatment. Findings related to that aspect of the study will be presented in a separate manuscript.

Remarkably, all vaccinated mice survived the acute phase of infection and progressed to a chronic disease state, which persisted for at least 10 weeks post-infection. During this period, lung bacterial burdens increased slowly but steadily in both sexes, reaching approximately 10⁸ CFU by week 10 (Fig. 1B).

H&E-stained lung sections revealed progressive granuloma formation, displaying similar histopathological patterns in both sexes (Fig. 2). By 6 weeks post-infection, lungs exhibited primarily cellular, non-necrotic lesions, which progressed to well-structured necrotic granulomas by week 8. These lesions continued to expand in size and complexity through week 10, consistent with the development of chronic pulmonary pathology. These granulomas were marked by central necrosis surrounded by dense immune cell infiltrates (Fig. 2).

**Figure 2.**
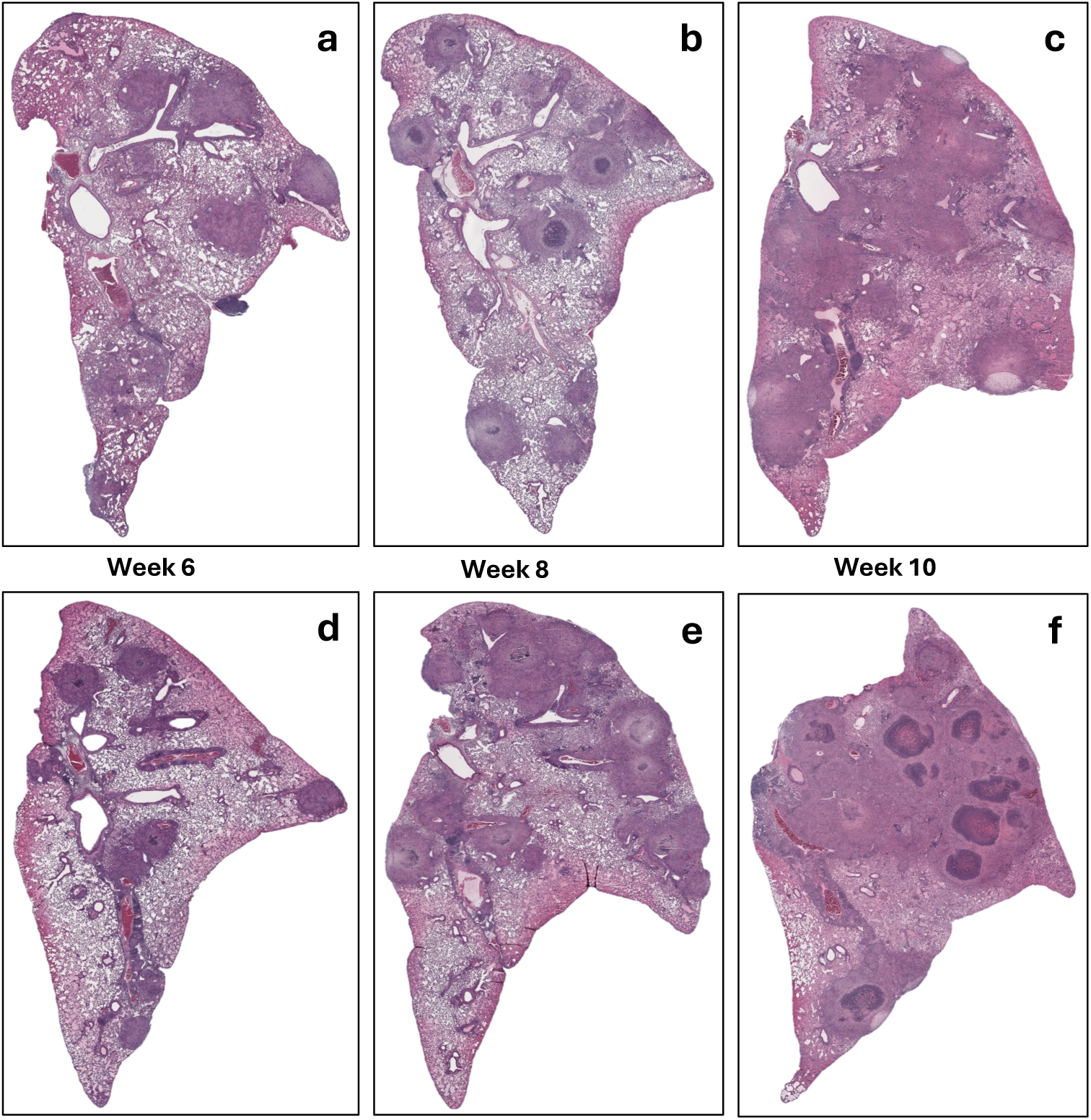
Histopathological Progression of Lung Lesions in Vaccinated Nos2^−/−^ Mice Infected with *Mtb* Erdman. Hematoxylin and eosin (H&E) staining of left lung sections from *Nos2*^−/−^ mice vaccinated with *Mtb* mc^2^6230 and challenged with *Mtb* Erdman. Panels show lesion progression with increasing structural complexity and necrosis at 6, 8, and 10 weeks post-infection: (a3c) female, (d3f) male mice. (Magnification 10×)

### Pimonidazole Staining Reveals Hypoxia Precedes Necrosis in TB Lesions

Hypoxia is a well-established hallmark of necrotic granuloma cores and is closely associated with advanced tuberculosis pathology [25]. To assess hypoxic regions within the lung, immunohistochemical staining for pimonidazole adducts, a hypoxia-specific probe, is commonly employed [26]. Because pimonidazole forms adducts only in viable hypoxic cells, the tissue distribution of these adducts provides an indication of hypoxic regions at the time of euthanasia.

To investigate the spatial and temporal dynamics of lesion hypoxia, immunohistochemical staining for pimonidazole adducts was performed on lung sections beginning at 4 weeks post-infection. At this early stage, pimonidazole adducts were detectable in the core of nascent granulomas, in cellular regions that had not yet progressed to necrosis. By week 8, staining became more spatially organized and was concentrated around the periphery of the developing necrotic core (Fig. 3), a pattern consistent with previous observations in necrotic granulomas of C3HeB/FeJ mice, where hypoxic staining forms a rim around necrotic regions [27]. These hypoxia profiles were consistent across both male and female *Nos2*-deficient mice in our study.

**Figure 3:**
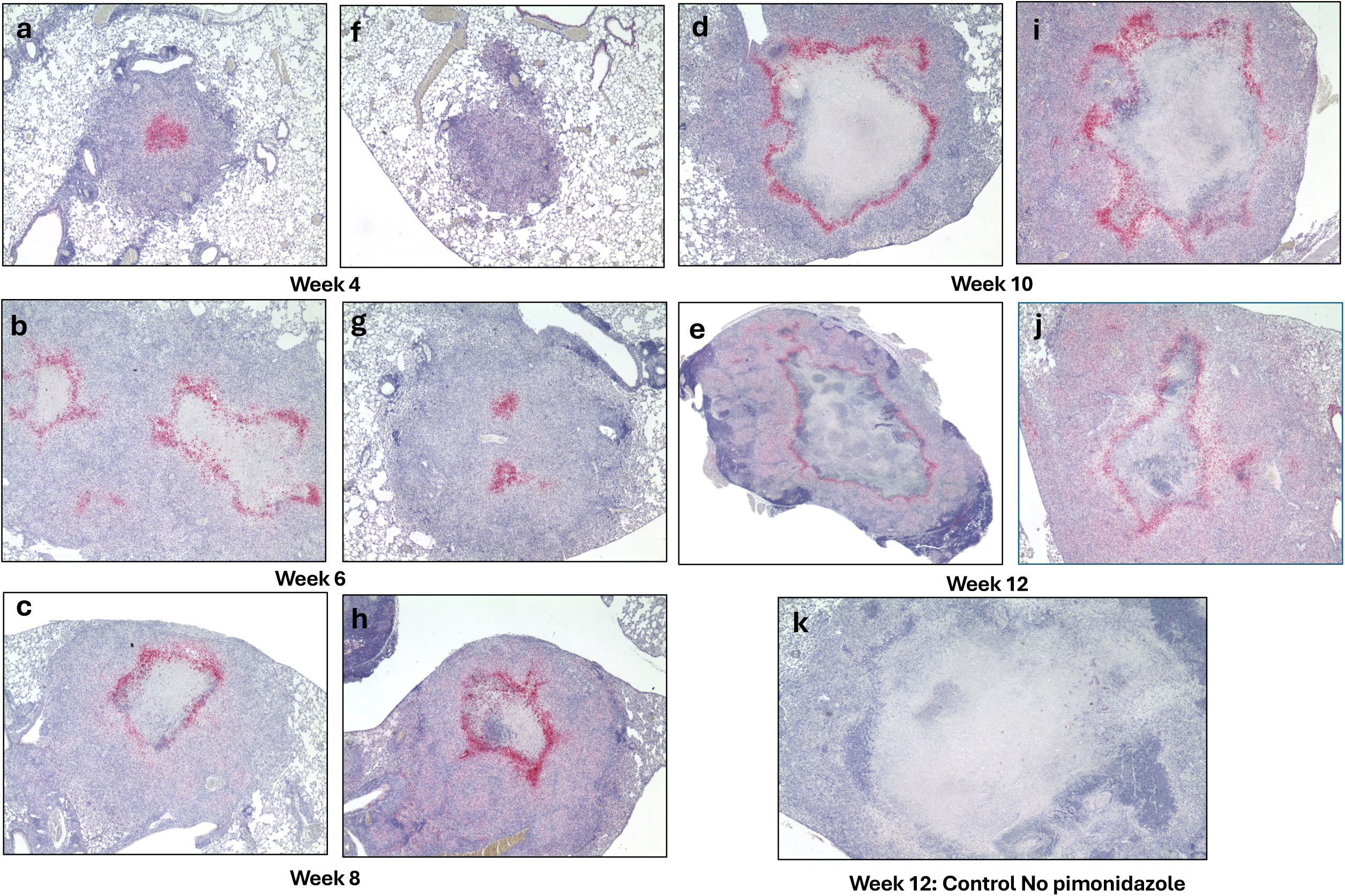
Detection of hypoxia in developing necrotic lung lesions of *Nos2*^−/−^ mice in the vaccinated model, using pimonidazole staining. Pimonidazole staining was used to detect hypoxia in lung lesions of vaccinated *Nos2*^−/−^ mice infected with *Mtb* Erdman. Time points at 4 (a and f), 6 (b and g), 8 (c and h), 10 (d and i), and 12 (e and j) weeks post-infection show progressive accumulation of pimonidazole adducts, indicating expanding hypoxic regions as maturing granuloma lesions form. Panels a3e show female mice; f3j show males and a week 12 infected mouse that did not receive pimonidazole served as a negative control to confirm staining specificity (k). (Magnification 40×)

Together, these findings indicate hypoxia precedes necrosis and likely contributes to structural evolution of necrotic granulomas in *Nos2*-deficient mice, and develops in regions consistent with patterns reported in other mouse models of hypoxic, necrotizing tuberculosis lesions [26, 28-30]

### Granuloma Microenvironment Contributes to Anti-Tubercular Drug Tolerance

To investigate how the granuloma microenvironment influences the efficacy of anti-tubercular drugs, including those used clinically to treat both drug-susceptible (DS-TB) and multidrug-resistant tuberculosis, we assessed the activity of multiple agents with distinct mechanisms of action in vaccinated *Nos2*-deficient mice (both male and female) at two distinct stages of infection. Drug treatment was initiated either: at week 6 post-infection, when granulomas were predominantly cellular and non-necrotic, and continued for 2 or 4 weeks; or at week 8 post-infection, when granulomas were well-structured, hypoxic, and necrotic, followed by a 2-week treatment course. Lung bacterial burdens were quantified at each post-treatment time point, and log_10_ CFU reductions were calculated relative to the corresponding pre-treatment baselines (week 6 or week 8, respectively).

Due to the lack of significant differences in CFU counts between male and female mice, data from both sexes were pooled and presented together for the analysis of drug activity (Fig. 4). As expected, when treatment was initiated at week 6 post-infection, untreated mice exhibited progressive infection, with lung bacterial burdens increasing by 0.56 log_10_ CFU by week 8 and a total increase of 0.85 log_10_ CFU by week 10. In contrast, drug-treated groups showed varying degrees of bacterial clearance.

**Figure 4.**
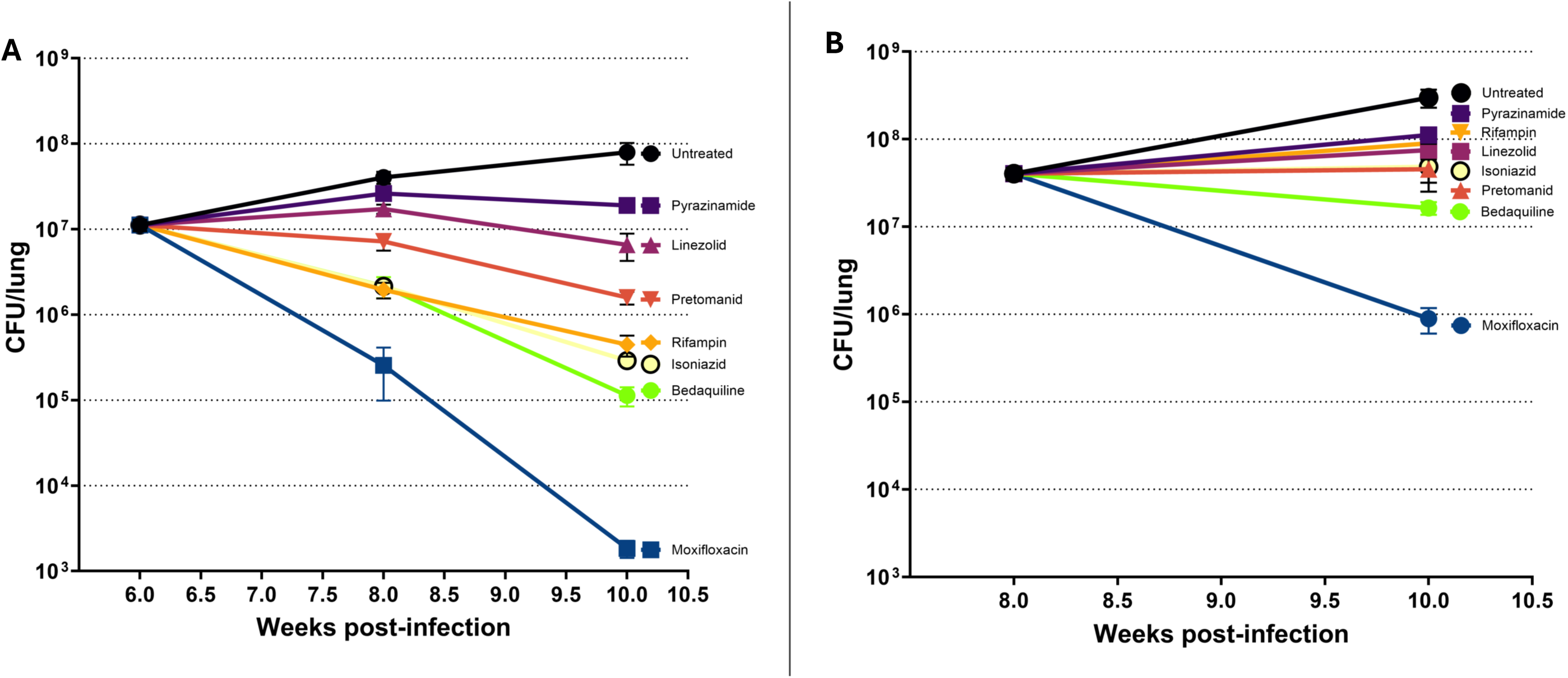
Influence of Lesion Structure on Drug Efficacy in Vaccinated *Nos2*^−/−^ Mice Infected with *Mtb*. Drug treatment was initiated either before (6 weeks post-infection, panel A) or after (8 weeks, panel B) the development of structured lung lesions, to assess the impact of lesion formation on drug efficacy responses. Drugs were administered by oral gavage 5 days per week. The following doses were used: Isoniazid 3 25 mg/kg, Pyrazinamide 3 150 mg/kg, Moxifloxacin 3 200 mg/kg, Rifampicin 3 10 mg/kg, Bedaquiline 3 25 mg/kg, Linezolid 3 100 mg/kg, and Pretomanid 3 75 mg/kg. Each data point represents the mean ± s.e.m. of data from 8310 mice per group (including both males and females).

Among the compounds tested, moxifloxacin (MXF, 200 mg/kg), a fluoroquinolone that targets DNA gyrase and topoisomerase IV, demonstrated the strongest bactericidal activity, with a 1.64 log_10_ CFU reduction by week 8, increasing to 3.78 log_10_ by week 10, indicating potent and sustained bacterial clearance. This result is consistent with the substantial bactericidal activity of MXF in stationary phase and hypoxia [31, 32], which may mimic attributes of this chronic infection, in which we observed significant hypoxic regions.

Both bedaquiline (BDQ, 25 mg/kg), an ATP synthase inhibitor, and isoniazid (INH, 25 mg/kg), a prodrug that inhibits mycolic acid synthesis, exhibited significant activity, with 0.72 log_10_ reductions at week 8, improving to 1.99 and 1.59 log_10_, respectively, by week 10. While BDQ has been shown to retain activity against *Mtb* under starvation and hypoxia, INH loses activity in all non-replicating persistence assays [1]. The efficacy of these compounds at early time points in our *Nos2* model may be an indication of the heterogenous microniches present in the model and replicates Early Bactericidal Activity (EBA) clinical results [33].

Rifampicin (RIF, 10 mg/kg), an RNA polymerase inhibitor, yielded a 1.4 log_10_ CFU reduction by week 10, consistent with its known bactericidal activity in DS-TB [34].

Pretomanid (PMD, 75 mg/kg), a nitroimidazole that releases reactive nitrogen species and inhibits mycolic acid biosynthesis; and linezolid (LZD, 100 mg/kg), a protein synthesis inhibitor targeting the 50S ribosomal subunit, produced more modest reductions, with CFU declines not exceeding 0.85 log_10_ by week 10. These results are consistent with the slower bactericidal activity of PMD and LZD in EBA studies [35, 36].

Pyrazinamide (PZA, 150 mg/kg) exhibited limited bacteriostatic effects, with a 0.37 log_10_ CFU increase by week 8 and 0.23 log_10_ increase by week 10. This is consistent with its known reduced potency in necrotic environments and negligible EBA [13, 27, 37, 38].

When treatment was initiated at week 8 post-infection, moxifloxacin demonstrated the most potent bactericidal activity, achieving a 1.66 log_10_ CFU reduction. This robust and sustained bacterial clearance is consistent with prior findings showing moxifloxacin’s superior sterilizing potential in chronic TB models [39, 40]. Bedaquiline exhibited moderate efficacy, producing a 0.39 log_10_ CFU decline, consistent with its delayed bactericidal kinetics previously reported in murine TB studies [41, 42]. However, several other drugs exhibited little to no further bactericidal activity during this treatment window. Pretomanid and isoniazid were associated with slight increases in bacterial burden (0.05 and 0.08 log_10_ CFU, respectively), suggesting limited efficacy under these conditions [42-44]. Linezolid and rifampin produced respective increases of 0.27 and 0.35 log_10_ CFU, indicating bacterial persistence as RIF accumulates in the caseum, and LZD has been shown to similarly lose efficacy in *Mtb*-infected C3HeB/FeJ mice developing caseous lesions [8, 13]. Pyrazinamide exhibited the least favorable outcome, with a 0.44 log_10_ CFU increase, this result parallels what has been observed in guinea pigs, which develop necrotic hypoxic granulomas that are refractory to pyrazinamide treatment [45]. In the absence of treatment mice displayed a 0.87 log_10_ CFU increase, consistent with ongoing disease progression in the absence of therapy.

### Isoniazid Reduces Granuloma Progression but Does Not Prevent Necrosis

To evaluate the effectiveness of INH monotherapy on granuloma lesion development in both male and female *Nos2*-deficient mice, lungs from untreated control animals were harvested in parallel with those from INH-treated mice at identical time points, and histopathological analysis was performed (Fig. 5). When the treatment started at 6 week post-infection, INH-treated lungs exhibited fewer and less mature granuloma lesions compared to untreated controls (Fig. 5A), indicating a delay in granuloma progression, consistent with previous findings demonstrating that INH reduces bacterial burden and early lesion formation [26, 46]. However, despite this attenuation, necrotic cores still developed in treated mice, particularly at later stages of infection (week 10) (Fig. 5B). This aligns with prior observations that INH monotherapy, while bactericidal against actively replicating *M. tuberculosis*, fails to eliminate non-replicating or hypoxic bacilli, which continue to drive inflammation and necrosis [25, 43, 47].

**Figure 5:**
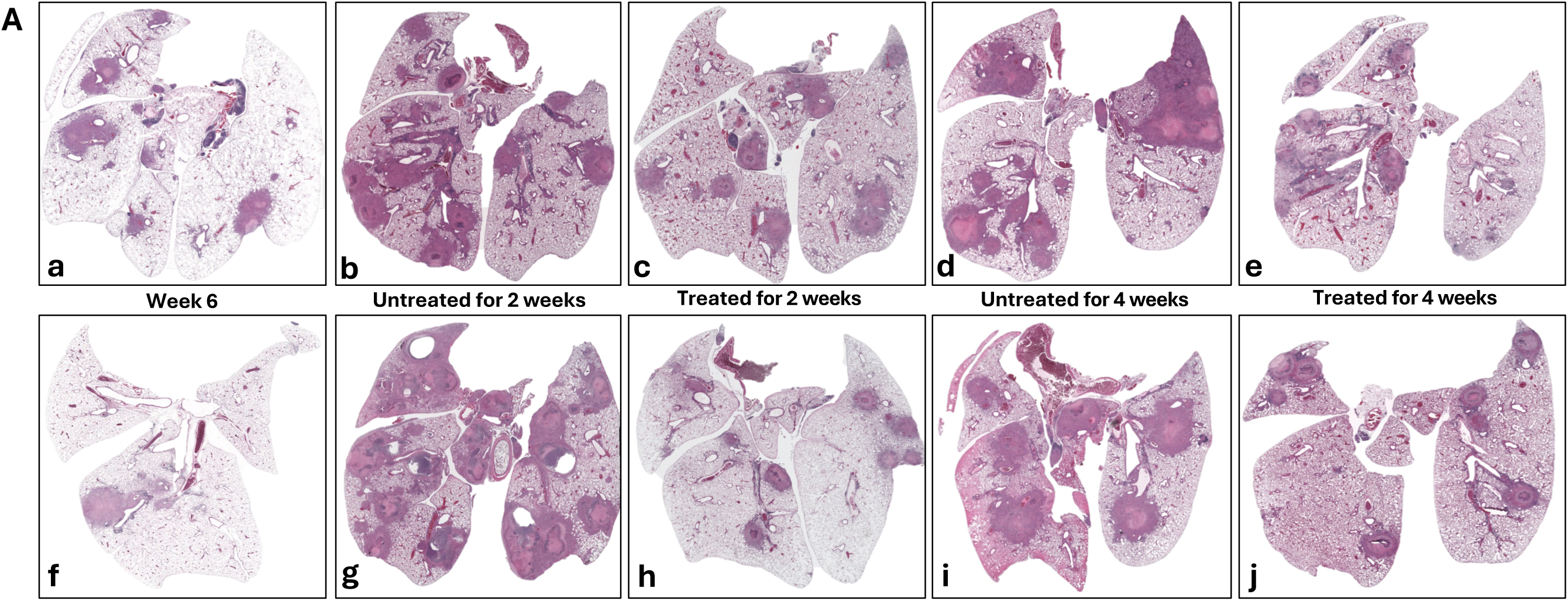
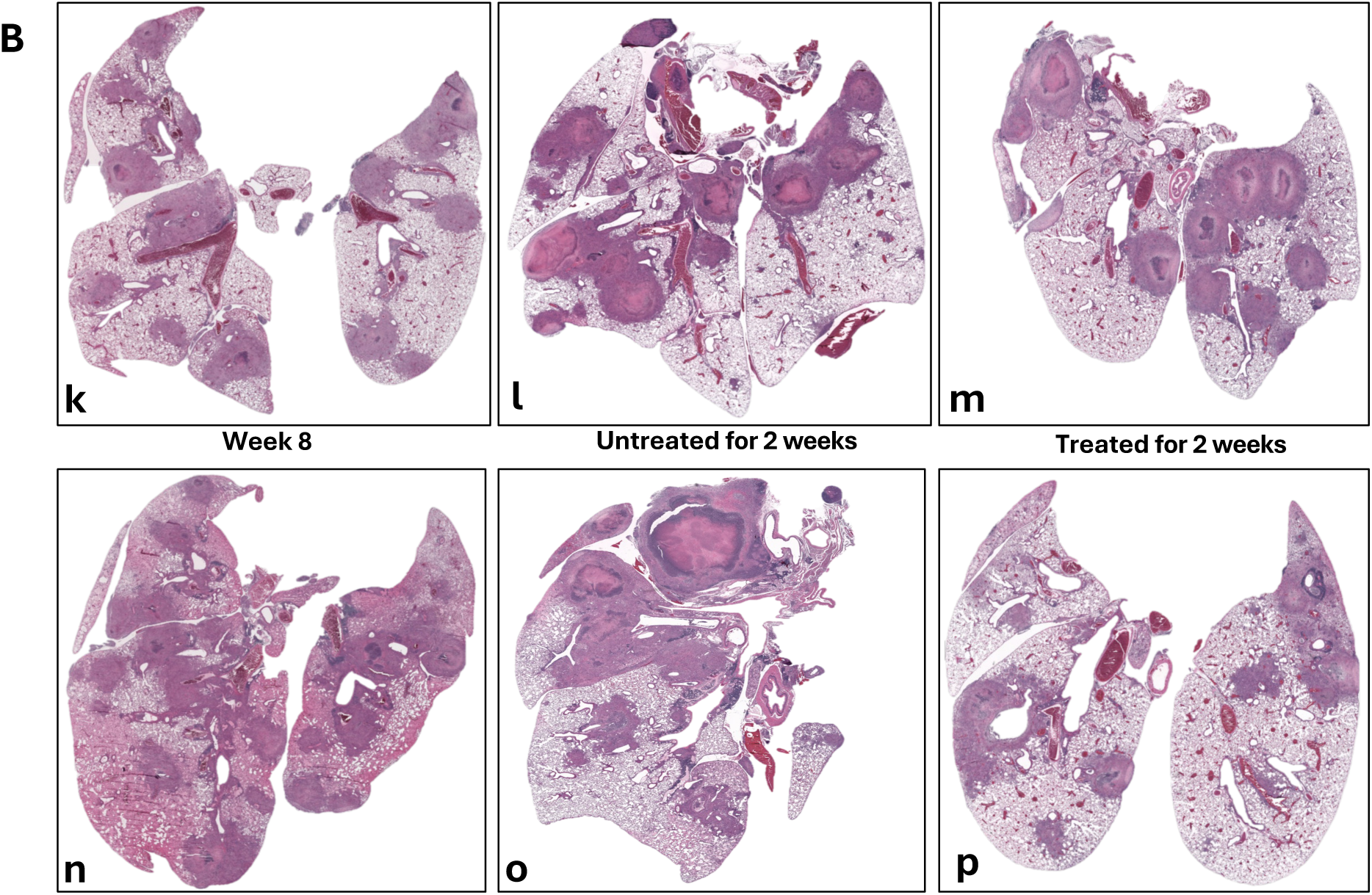
Impact of isoniazid (INH) treatment on lung tissue in vaccinated *Nos2*^−/−^ mice infected with *Mtb* Erdman. Histological analysis of H&E-stained lung sections from a representative vaccinated *Nos2^{/{^* mouse treated with INH (25 mg/kg), starting either at 6 weeks (Panel A) or 8 weeks (Panel B) post-infection. Early treatment leads to improved lesion resolution, whereas delayed treatment is less impactful due to the establishment of mature granuloma formation. (Magnification: 10×) **Panel A** illustrates lung pathology in female (a3e) and male (f3j) mice following initiation of INH treatment at week 6 post-infection. (a, f): Baseline prior to treatment initiation (week 6); (b, g): Two weeks without treatment (week 8, untreated); (c, h): Two weeks with INH treatment (week 8, treated); (d, i); Four weeks without treatment (week 10, untreated) and (e, j): Four weeks with continued INH treatment (week 10, treated) **Panel B** shows lung pathology in female (k3m) and male (n3p) mice when INH treatment start at week 8 post-infection. (k, n): Baseline prior to treatment initiation (week 8), (l, o): Two weeks without treatment (untreated); (m, p): Two weeks with INH treatment (treated).

## Discussion

This study establishes a *Nos2*-deficient mouse model of progressive, necrotizing tuberculosis, offering insights into the temporal dynamics of granuloma maturation, lesion hypoxia, and drug efficacy in a chronic, high-burden host environment. Building upon earlier findings where *Nos2*-deficient mice were reported to begin to slowly clear an infection with *Mtb* R1Rv from 3 weeks post-infection [19], we demonstrate that infection with the partially attenuated *Mtb* R1Rv strain leads to a progressive, high-burden infection in *Nos2*-deficient mice over a 10-week period (Fig. 1A). This subtle divergence suggests that disease progression in this model may be influenced by inoculum dose, which was slightly higher in our model to provide a higher burden to better evaluate highly efficacious treatments. Notably, both male and female mice developed similar lung bacterial burdens, indicating minimal sex-based differences in disease outcome. Histological analysis confirmed the formation of well-structured granulomas with necrotic centers and lesion hypoxia (Supplementary Fig. 1), consistent with advanced disease comparable to human tuberculosis and other susceptible mouse strains such as C3HeB/FeJ [27, 48].

Following aerosol Erdman challenge in mc² 6230–vaccinated *Nos2*-deficient mice, a similar pathological course was observed: steady bacterial growth reaching ∼10⁸ CFU by week 10, along with the development of necrotic granulomas in both sexes (Fig. 1B). Importantly, vaccinated mice survived well beyond the acute phase of the disease, highlighting a degree of protective efficacy even in the absence of iNOS. Although nitric oxide (produced via Nos2/iNOS) has long been associated with antimicrobial activity, recent evidence suggests that its protective role during *Mtb* infection lies not primarily in direct bacterial killing, but in modulating the host inflammatory response by suppressing a pathological over-recruitment of neutrophils, which leads to necrosis and enhanced bacterial proliferation [49]. Our finding therefore suggests that vaccine-induced adaptive immunity can delay disease progression, even in the absence of innate effector mechanisms such as nitric oxide production.

In comparison, while C3HeB/FeJ mice exhibit considerably more variable disease kinetics and pathology, mortality in this model can fluctuate depending on the infectious dose and strain virulence. High bacterial burden is not consistently observed across animals; instead, this strain is characterized by pronounced inter-mouse heterogeneity. Some mice experience delayed progression, whereas others rapidly develop fulminant disease characterized by extensive necrosis and tissue liquefaction [12, 13, 26]. This variability contrasts sharply with the more uniform disease course seen in *Nos2*-deficient mice, which exhibit more consistent granuloma structure and progression. As such, the *Nos2*-deficient model complements alternative models and offers a robust and reproducible platform for evaluating novel therapeutic agents or optimizing current treatment regimens.

Immunohistochemical analysis to detect pimonidazole adducts revealed that hypoxia was present within nascent granulomas as early as week 4 post-infection, and became increasingly organized around necrotic cores by week 8 (Fig. 3), a pattern consistent with observations in other necrotizing TB animal models [28, 50, 51]. This temporal progression strongly supports the hypothesis that hypoxia is not merely a consequence of necrosis but rather precedes it and may actively contribute to its development. These findings align with previous studies in C3HeB/FeJ mice and human TB granulomas, where oxygen depletion has been implicated as a driver of necrosis resulting from impaired vascularization and inflammation-induced tissue remodeling [25, 26, 51]. The early presence of hypoxic, non-necrotic lesions provides a potential window for therapeutic intervention before irreversible structural damage occurs. Moreover, the consistent presence of hypoxia across sexes and challenge strains highlights the generalizability of this phenomenon within the model.

The vaccinated mouse model revealed pronounced differences in drug efficacy depending on the timing of treatment initiation and the granuloma stage. When therapy began at week 6, before widespread necrosis, multiple first- and second-line agents demonstrated strong bactericidal activity. Moxifloxacin showed the greatest efficacy, followed by bedaquiline, isoniazid, and rifampicin, consistent with their known mechanisms targeting actively replicating *Mtb* populations (Fig. 4A) [43, 52, 53]. However, when treatment began at week 8, after lesions had become necrotic, drug efficacy was substantially reduced (Fig. 4B). Only moxifloxacin retained appreciable activity, while most other agents, including bedaquiline, isoniazid, linezolid, and pretomanid, showed minimal or no net reduction in CFU; some even allowed modest increases in bacterial burden. Pyrazinamide exhibited the least activity under these conditions, as has been observed in C3HeB/FeJ mice and that others have attributed to the neutral pH of caseum [12, 13, 54, 55].

The reduced activity of several drugs in necrotic lesions, even where they were effective in early-stage cellular granulomas, further validates the importance of lesion-specific pharmacodynamics. Notably, moxifloxacin’s retained efficacy regardless of lesion state supports its potential for sterilizing challenging microenvironments and supporting its inclusion in short-course regimens for drug-resistant TB [52, 56]. The limited efficacy of pyrazinamide and pretomanid in these conditions align closely with prior findings by Gengenbacher et al. (2017) and Irwin et al. (2016), raising critical concerns about their utility in late-stage TB unless combined with agents targeting persistent bacterial populations [12, 28]. The observation of substantial efficacy of most drugs when treatment was initiated at 6 weeks post infection, when hypoxia has already developed, brings into question the relative importance of hypoxia as a drug tolerance-inducing condition. Indeed, the disappointing efficacy of the anaerobic-activated drug metronidazole in a clinical trial suggests other environmental conditions may be more important drivers of persistence [57].

Gengenbacher et al. further emphasized that drug activity in their *Nos2*-deficient model correlates strongly with human clinical outcomes, and that treatment failure in hypoxic lesions may be more attributable to pathology-driven drug tolerance rather than genetic resistance mechanisms [28]. These insights are fully consistent with effects we observed in advanced disease, reinforcing the notion that effective late-stage TB therapy may require combining existing drugs with new agents able to overcome microenvironment-driven tolerance.

Overall, these findings highlight key barriers to drug efficacy in necrotic granulomas, drugs that are effective in early-stage disease may fail in mature necrotic lesions due to: impaired drug penetration into necrotic caseum, altered host metabolism, and the presence of hypoxic, non-replicating bacilli, all of which contribute to intrinsic drug tolerance in advanced lesions [8, 58]. This insight has important implications for both preclinical drug evaluation and clinical regimen design, emphasizing the need to consider lesion stage when developing effective TB therapies.

Histopathological evaluation of lung tissue in INH-treated mice revealed a delay in granuloma progression, with fewer and less mature lesions when therapy was initiated early. Nevertheless, necrosis continued to develop, consistent with INH’s inability to sterilize non-replicating bacilli (Fig. 5). This outcome is consistent with the INH’s narrow spectrum of activity, which targets actively replicating *Mtb* while leaving behind persistent subpopulations that contribute to ongoing inflammation and driving lesion progression. These findings reinforce the inadequacy of monotherapy within complex lesion environments and highlight the critical need for multidrug regimens capable of sterilizing heterogeneous bacterial populations across diverse pathological niches.

## Materials and Methods

### Ethics Statement and Mouse Strain

Mice were bred and housed in the Trudeau Institute’s barrier-specific pathogen-free (SPF) animal facilities. Animals were maintained on UV/particulate-filtered, acidified water (pH 2.5–3.0) and autoclaved LabDiet 5P04 rodent chow, housed in sterile individually ventilated cages (IVCs) with sterile beta chip hardwood bedding and sterile enrichment. Trudeau Institute’s animal facilities are accredited by the Association for Assessment and Accreditation of Laboratory Animal Care International (AAALAC), licensed by the USDA, and maintain an Animal Welfare Assurance on file with the National Institutes of Health (NIH) Office of Laboratory Animal Welfare.

All procedures were conducted in accordance with protocols approved by the Institutional Animal Care and Use Committee (IACUC), including experimental design, animal welfare monitoring, humane endpoints, and use of AVMA approved euthanasia methods. Animal care protocols, housing conditions, and monitoring procedures were consistent across both biosafety level II and III facilities. Mice were maintained under sterile conditions with standard housing, bedding, food, and water. Animal health was monitored regularly, including monthly weight measurements. Humane endpoints were defined as a body weight loss exceeding 20% of the pre-experimental weight, accompanied by clinical signs of distress such as hunched posture, inactivity, isolation, or greasy, severely ruffled fur.

The *Nos2*-deficient mouse strain B6.129P2-*Nos2^tm1Lau^*/J (strain #002609), aged 4–6 weeks were bred in-house from stock obtained from The Jackson Laboratory (Bar Harbor, ME). Mice were initially housed in a biosafety level II (BSL-2) facility, where they received subcutaneous vaccination with *Mycobacterium bovis* strain BCG Pasteur (TMC#1011, from the Trudeau Mycobacterial Culture Collection, Trudeau Institute, Saranac Lake, NY) or the attenuated *Mtb* strain mc^2^6230. They were subsequently transferred to a biosafety level III (BSL-3) facility for aerosol infection with the virulent *Mtb* H37Rv or Erdman pYUB1785 strain or, in naïve mice, with the *Mtb* strain R1Rv.

### Bacterial growth conditions and aerosol infections

#### Naïve model: *Mycobacterium tuberculosis* R1Rv Infection

The partially attenuated *Mtb* strain R1Rv (TMC#205) was cultured in complete 7H9-T medium. This medium consisted of Middlebrook 7H9 base (BD Difco, Franklin Lakes, NJ) supplemented with 10% (v/v) oleic acid-albumin-dextrose-catalase (OADC), prepared from oleic acid (0.5 g/L, Thermo Fisher Scientific, Waltham, MA), albumin (50 g/L, Fisher Scientific, Hampton, NH), dextrose (20 g/L, Fisher Scientific), and catalase (0.04 g/L, Thermo Fisher Scientific). The medium also contained sodium chloride (8 g/L, Fisher Scientific), glycerol (0.5% v/v, Fisher Scientific), supplemented with 0.05% tyloxapol (Thermo Fisher Scientific).

Both sexes of *Nos2*-deficient mice, aged 6–8 weeks, were infected by aerosol exposure with approximately 1000 colony-forming units (CFUs) of *Mtb* R1Rv using a Glas-Col inhalation exposure system [59].

Mice were euthanized via CO₂ inhalation at predetermined post-infection time points. Whole lungs were aseptically removed and homogenized for viable bacterial count (CFU enumeration). In a separate aerosol challenge experiment, the left lung lobe was perfused and embedded for histological analysis as documented in the supplementary figures, while the remaining lobes were homogenized for CFU enumeration.

#### Vaccinated model: Vaccination with Attenuated *Mtb* mc² 6230 and Aerosol Challenge with Erdman-pYUB1785

For this model, the attenuated *Mtb* strain mc²6230 (kind gift of Dr. William R. Jacobs, Jr., Albert Einstein College of Medicine, Bronx, NY) was cultured in complete 7H9-T medium supplemented with 24 µg/mL D-pantothenate. Ten milliliter cultures were grown in 30 mL inkwell bottles at 37 °C with constant shaking at 150 rpm until reaching mid-log phase (OD₆₀₀ = 0.4–0.8). The culture was then diluted to an OD₆₀₀ of 0.4 and 100 µL was administered subcutaneously into mice.

Four weeks post-vaccination, mice were challenged with the virulent *Mtb* Erdman-pYUB1785 strain.

To construct the stable persister reporter strain used for challenge of vaccinated mice, *Mtb* Erdman was co-electroporated with two plasmids: pYUB1785, a *pdnaK*::tdTomato pL5::mVenus construct [24] introduced into a vector bearing L5attP integration site but lacking integrase; and pYUB1505, a suicide vector expressing integrase. The resulting strain, *Mtb* Erdman-pYUB1785 strain was grown in complete 7H9-T medium supplemented with kanamycin (50 µg/mL) to an OD_₆₀₀_ of 0.4–0.6, and the inoculum was diluted to an OD_₆₀₀_ of 0.02 for aerosol infection. Approximately 100 CFU were delivered via aerosol using a Glas-Col inhalation exposure system, as described above.

To determine the initial bacterial implantation in the lungs, five mice were sacrificed one day following aerosol exposure in each infection model.

#### Vaccinated model: Drug Preparation and In Vivo Treatment

For drug treatment studies, mice received individual anti-tuberculosis drugs administered as monotherapy. Isoniazid (INH), Pyrazinamide (PZA), Moxifloxacin (MXF), and Linezolid (LZD) were purchased from Acros Organics; Rifampicin (RIF) was obtained from Sigma-Aldrich, and Bedaquiline (BDQ) and Pretomanid (PMD) were purchased from MedChemExpress. Drug formulations were prepared fresh weekly as follows: INH at 25 mg/kg, PZA at 150 mg/kg, and MXF at 200 mg/kg were dissolved in sterile water. RIF at 10 mg/kg was first dissolved in 100% DMSO and then diluted in sterile water to achieve a final DMSO concentration of 5%. BDQ at 25 mg/kg was formulated in acidified 20% hydroxypropyl-β-cyclodextrin (HPβCD). LZD at 100 mg/kg was formulated in a mixture of 5% PEG 200 and 95% of 0.5% methylcellulose. PMD at 75 mg/kg was prepared in cyclodextrin micelle (CM-2) formulation, containing 10% HPβCD (Sigma) and 10% lecithin (ICN Pharmaceuticals Inc., Aurora, OH). All suspensions were sonicated or rotated overnight at 4 °C to ensure uniformity.

Treatment began either 6- or 8-weeks post-infection, depending on the experimental group. Drugs were administered via oral gavage at a volume of 200 μL per mouse, 5 days per week and mice treated with sterile water served as negative controls. At least four mice from each group were euthanized via CO₂ inhalation after 2 and 4 weeks of treatment for analysis.

### Viable Count of Bacteria in Lungs

On the designated harvest day, lungs were aseptically removed from each mouse and placed in sterile glass homogenizer tubes containing 4.5 mL of 0.85% saline, and the tissues were homogenized using a mechanical homogenizer (Glas-Col Inc., Terre Haute, IN). To quantify the viable bacterial burden, lung homogenates were serially diluted in 0.85% saline and plated on Middlebrook 7H11 agar supplemented with OADC and glycerol. For samples from mice treated with BDQ, homogenates were plated on 7H11 agar containing 0.4% activated charcoal to prevent drug carry-over. Plates were incubated at 37 °C + 5% CO₂, and CFU were enumerated after a minimum of 21 days. For charcoal-containing plates, longer incubation times were applied as needed to allow visible colony development.

### Histopathology and Pimonidazole Immunohistochemical Staining

Beginning at 4 weeks post-aerosol infection, lungs were inflated and fixed in 4% methanol-free formaldehyde in phosphate-buffered saline (PBS). Fixed tissues were processed, embedded in paraffin, and sectioned 5 microns thick for histological analysis. Tissue sections were mounted on glass slides, deparaffinized, and stained with hematoxylin and eosin (H&E). Whole-slide scans were obtained using the Microscopes International uSCOPE HXII slide scanner with a 10× objective and uSCOPE Navigator 4 software. To visualize hypoxic regions in lung tissues, pimonidazole immunohistochemical staining was performed using the Hypoxyprobe Omni kit (Hypoxyprobe Inc., Burlington Massachusetts). Mice were administered 100 mg/kg pimonidazole hydrochloride intraperitoneally at least 1.5 hours before sacrifice. Lungs from pimonidazole-treated and untreated control animals were instilled and immersed in 4% methanol-free formaldehyde for a minimum of 24 hours. Tissues were then processed into paraffin blocks using standard procedures.

Paraffin sections (5 μm) were mounted on charged slides, dried, and deparaffinized through successive xylenes and decreasing graded ethanol washes before rehydration in distilled water. Heat-induced epitope retrieval was performed by incubating slides in citrate buffer using a steamer for 40 minutes, followed by cooling and rinsing in PBS. Endogenous peroxidase and phosphatase activity was blocked using BLOXALL solution (Vector Laboratories), and non-specific binding was minimized with 5% normal horse serum. Slides were then incubated with rabbit anti-pimonidazole primary antibody, followed by washing in PBS and detection using an anti-rabbit alkaline phosphatase secondary kit (Vector Laboratories). The hypoxic regions were visualized using Vector Red alkaline phosphatase substrate (Vector Laboratories). Tissues from mice that have not received pimonidazole served as negative controls to monitor for non-specific substrate development. All sections were counterstained with hematoxylin and scanned at high resolution using a Nikon Eclipse 50i brightfield microscope equipped with a SPOT Imaging CMOS camera and corresponding imaging software.

## Acknowledgements

The attenuated *Mtb* strain mc^2^ 6230 was a kind gift of Dr. William R. Jacobs Jr. of Albert Einstein College of Medicine. The plasmids pYUB1785 and pYUB1505 were the kind gifts of Dr. Paras Jain and Dr. William R. Jacobs Jr. of Albert Einstein College of Medicine. We are grateful for the efforts of Amanda Schneck and the animal care staff of Trudeau Institute, without whom this work would not have been possible. This research was supported by Trudeau Institute and the National Institute of Allergy and Infectious Diseases of the National Institutes of Health under Award Number R21AI159731. The content is solely the responsibility of the authors and does not necessarily represent the official views of the National Institutes of Health.

